# Age-related learning and working memory impairment in the common marmoset

**DOI:** 10.1101/2022.06.07.495172

**Authors:** Courtney Glavis-Bloom, Casey R Vanderlip, John H Reynolds

## Abstract

Aging is the greatest risk factor for the development of neurodegenerative diseases, yet we still do not understand how the aging process leads to pathological vulnerability. The research community has relied heavily on mouse models, but the considerable anatomical, physiological, and cognitive differences between mice and humans limit their translational relevance. Ultimately, these barriers necessitate the development of novel aging models. As a non-human primate, the common marmoset (*Callithrix jacchus*) shares many features in common with humans and yet has a significantly shorter lifespan (10 years) than other primates, making it ideally suited to longitudinal studies of aging. Our objective was to evaluate the marmoset as a model of age-related cognitive impairment. To do this, we utilized the Delayed Recognition Span Task (DRST) to characterize age-related changes in working memory capacity in a cohort of sixteen marmosets varying in age from young adult to geriatric. These monkeys performed thousands of trials over periods of time ranging up to 50 percent of their adult lifespan. To our knowledge, this represents the most thorough cognitive profiling of any marmoset aging study conducted to-date. By analyzing individual learning curves, we found that aged animals exhibited delayed onset of learning, slowed learning rate after onset, and decreased asymptotic working memory performance. These findings are not accounted for by age-related impairments in motor speed and motivation. This work firmly establishes the marmoset as a model of age-related cognitive impairment.

**Significance Statement:** Understanding the normal aging process is fundamental to identifying therapeutics for neurodegenerative diseases for which aging is the biggest risk factor. Historically, the aging field has relied on animal models that differ markedly from humans, constraining translatability. Here, we firmly establish a short-lived non-human primate, the common marmoset, as a key model of age-related cognitive impairment. We demonstrate, through continuous testing over a substantial portion of the adult marmoset lifespan, that aging is associated with both impaired learning and working memory capacity, unaccounted for by age-related changes in motor speed and motivation. Characterizing individual cognitive aging trajectories reveals inherent heterogeneity, which could lead to earlier identification of the onset of impairment, and extended timelines during which therapeutics are effective.

## INTRODUCTION

Age-related cognitive impairment is well-documented in rodents, non-human primates (NHPs), and humans across multiple domains including attention, executive function, and working memory (Glisky, 2007). Of these, working memory is particularly vulnerable to the aging process with impairment evident early on (Belleville et al., 1998; Voytko and Tinkler, 2004; Gazzaley et al., 2005; Cappell et al., 2010; Klencklen et al., 2017). Intact working memory is dependent on the dorsolateral prefrontal cortex (dlPFC) and hippocampus, the two brain areas that undergo the earliest morphological and functional changes with age (Rusinek et al., 2003; Morrison and Baxter, 2012).

Translationally relevant model systems are critical for understanding these neurobiological and cognitive changes with age in humans. While mouse models are important, their translational relevance is limited by considerable differences with human neuroanatomy, physiology, and cognition (Preuss, 1995; Izpisua Belmonte et al., 2015). NHPs are more similar to humans in each of these areas and are genetically more closely related to humans. Importantly, unlike rodents, NHPs have a clearly defined dlPFC, making them ideal models in which to investigate prefrontal-dependent cognitive functions, including working memory (Wise, 2008). They also exhibit age-related cognitive decline that closely parallels that of humans (Upright and Baxter, 2021).

Several groups have investigated the effects of age on working memory performance in macaque monkeys, with mixed conclusions. Some show that aged macaques require more experience than young to reach a learning criterion on the delayed non-match-to-sample (DNMS) task (Rapp and Amaral, 1989; Herndon et al., 1997; Comrie et al., 2018) and the delayed response task (Bachevalier et al., 1991; Voytko and Tinkler, 2004). Others, however, show that age-related impairment is more heterogeneous, with nearly half of middle-aged (20-24 years) and aged (25-31 years) macaques demonstrating working memory impairments, while the other half did not (Moss et al., 2007). Similar heterogeneity amongst the aging human population is also well-described (Nyberg et al., 2020). Understanding what drives this variability is critical to identifying what causes some individuals to sustain age-related cognitive impairment, while others remain unaffected.

Longitudinal study designs provide an essential platform to investigate variability in cognitive aging trajectories and can pinpoint the onset of impairment (Hofer and Sliwinski, 2001; McQuail et al., 2021). However, the long lifespans of macaque monkeys (25-40 years) severely limit their use in longitudinal aging studies. The common marmoset (*Callithrix jacchus*) is advantageous as a NHP model for studying age-related cognitive impairment because they mature quickly, are considered aged at 7-8 years old, and have the shortest lifespan of any anthropoid primate, typically living only 9-10 years (Abbott et al., 2003; Schultz-Darken et al., 2016; Tardif, 2019). It is thus feasible to study aging longitudinally in the marmoset.

The use of marmosets in neuroscience research has accelerated (Abbott et al., 2003), but only a handful of groups have investigated age-related cognitive impairment (Munger et al., 2017; Phillips et al., 2019; Sadoun et al., 2019; De Castro and Girard, 2021), with only one utilizing a longitudinal design (Rothwell et al., 2022), and in that study, working memory was not assessed.

In the present study, we assessed marmosets’ performance on the Delayed Recognition Span Task (DRST) to measure working memory throughout their lifespans. The DRST depends critically on the two brain areas that undergo the earliest morphological and functional changes with age: dlPFC and hippocampus (Beason-Held et al., 1999; Bor et al., 2006; Jeneson et al., 2010). Additionally, older humans (55-87 years old) and aged macaque monkeys (20-27 years) each perform the DRST more poorly, on average, than young adults (humans: 18-25 years old; macaques 5-10 years old) (Herndon et al., 1997; Moss et al., 1997; Maylor et al., 2006; Belham et al., 2013; Mazurek et al., 2015; Satler et al., 2015). Here we find that increasing age is associated with delayed acquisition of task rules, slowed rate of learning once the rules have been established, and impaired working memory.

## MATERIALS AND METHODS

### Subjects

Sixteen (eight male, eight female) common marmosets (*Callithrix jacchus*) were trained to use touch screen computers installed in their home cages. Marmosets were between 1.7 and 14.6 years of age at the beginning of training and between 3.2 and 15.6 years at the conclusion of the study (Figure 1). Each of these 16 marmosets was tested on a working memory task, and a subset of 12 were also tested on tasks to measure motor function and motivation. All marmosets were tested concurrently for 1-3 hours on 3-5 days per week. No food or water restriction was used. Marmosets were singly or pair housed in cages with visual and auditory access to other marmosets in cages that contained a variety of enrichment items including hammocks and manzanita branches. Paired marmosets were separated during cognitive testing and were acclimated to separation prior to commencement of testing to minimize stress. All experiments were conducted in compliance with the Institutional Animal Care and Use Committee of the Salk Institute for Biological Studies and conformed to NIH guidelines.

**Figure 1.**
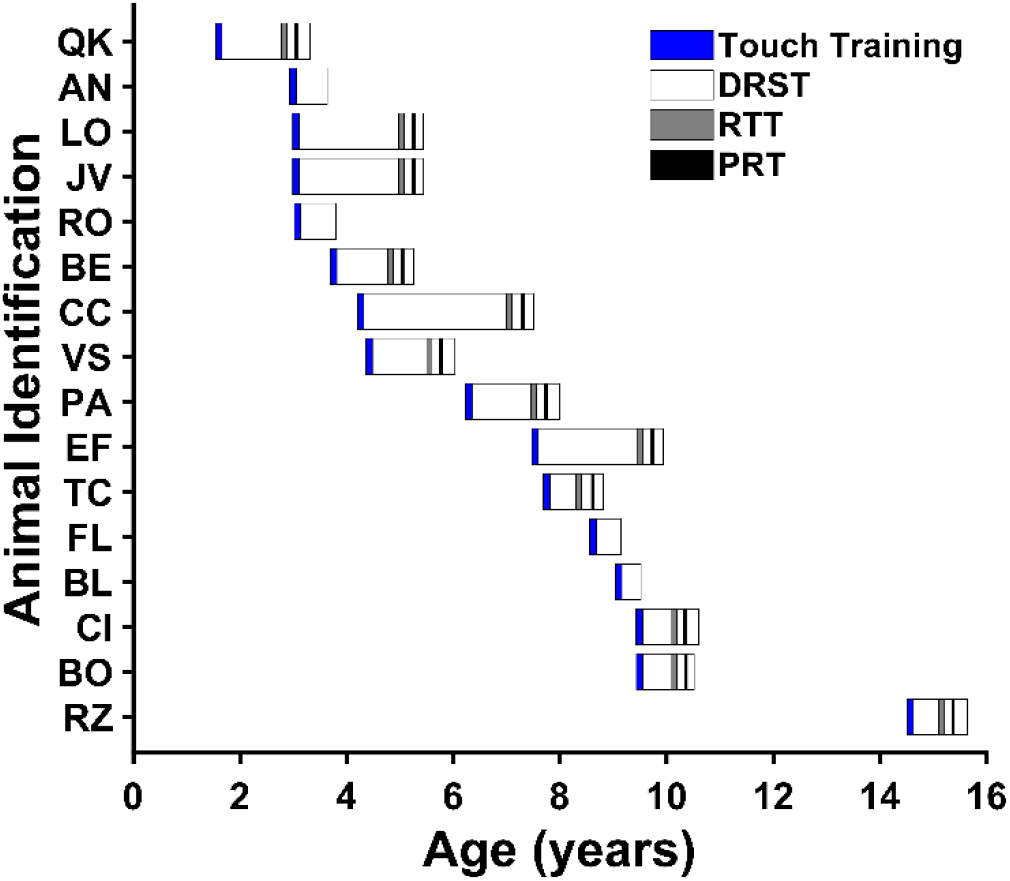
Cognitive testing across the marmoset lifespan. To enable assessment of age-related differences in acquisition of the Delayed Recognition Span Task (DRST), marmosets began training at ages that conjointly covered the lifespan. The age at which each marmoset learned to touch stimuli on the screen to earn juice rewards (Touch Training) is indicated in blue. The duration of longitudinal DRST testing is indicated for each marmoset in white. Age at time of control task testing is indicated in gray (Reaction Time Task for motor speed, RTT) and black (Progressive Ratio Task for motivation, PRT).

### Equipment

#### Cages

Marmoset home cages were modified to include a testing chamber in one of the upper corners of the cage, accessed via a small doorway (Figure 2C-D). During cognitive testing, the doorway was open, and marmosets could freely enter and exit the chamber. The doorway was closed at all other times, preventing marmosets from accessing the chamber.

**Figure 2.**
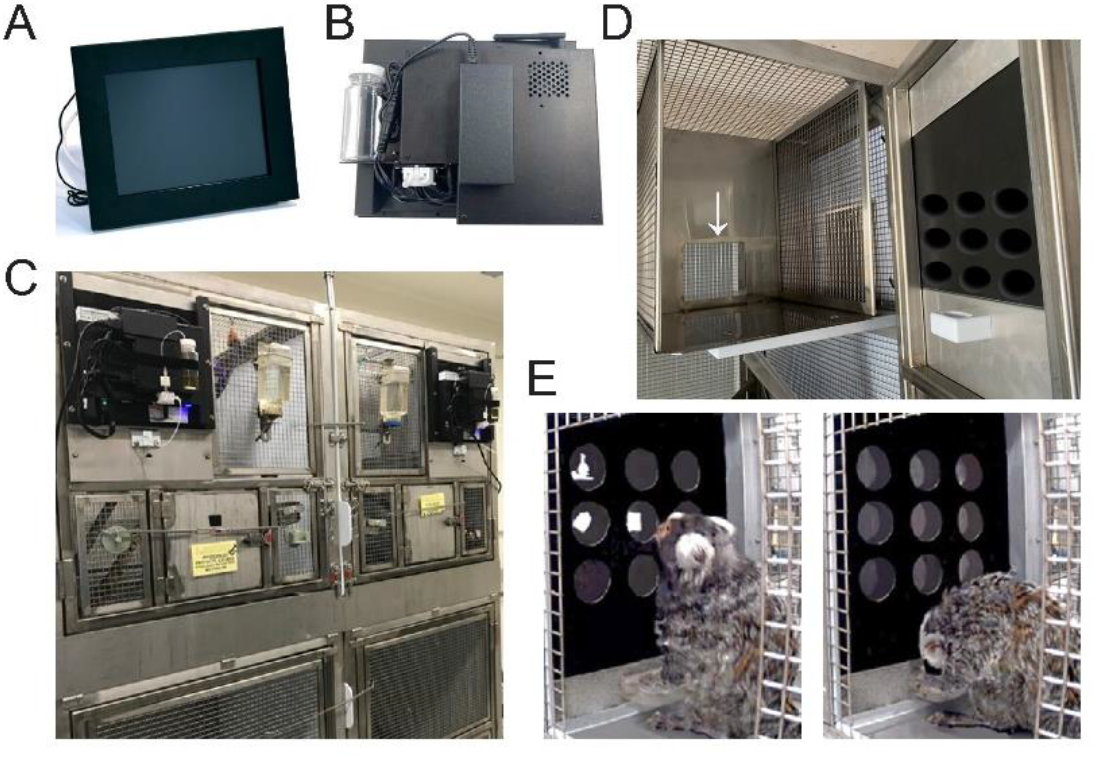
Equipment used for home cage cognitive testing. A & B) Touchscreen stations containing a 10.4-inch infrared screen and integrated peristaltic pump for dispensing liquid rewards were C) mounted on the front of each marmoset home cage and D) accessed via a small doorway (white arrow) to a testing chamber built into the home cage. E) A plastic screen covering (‘mask’) with nine equally sized and spaced holes (diameter = 1.5 in) arranged in three rows of three, was positioned in front of the screen. These holes provided access to the nine potential locations where stimuli could appear during cognitive testing.

#### Touchscreen stations

Touchscreen stations (Figure 2A-B; Lafayette Instrument Company, Lafayette, IN) were mounted on the front of the marmoset home cage (Figure 2C) and were accessible from the testing chamber built into the cage (Figure 2D). Each system contained a 10-inch infrared touchscreen with integrated peristaltic pump for dispensing of liquid rewards into a “sink” below the screen (Figure 2D-E). The infrared technology enabled highly reliable and sensitive detection of marmoset touches. A plastic screen covering (termed a ‘mask’) with nine equally sized and spaced holes (diameter = 1.5 in) arranged in three rows of three, was positioned in front of the screen (Figure 2E). These holes delineated the potential locations in which stimuli could appear during cognitive testing.

#### Software

Cognitive tasks were programmed using Animal Behavior Environment Test (ABET) Cognition software (Lafayette Instrument Company, Lafayette, IN) that controlled all aspects of the task including the number and order of trials, timing, stimuli selection and display location, and delivery of liquid rewards. The software also recorded several measures of task performance including trial number, correct and incorrect responses, the location on the screen of the correct choice and the chosen location, and response latency. Stimuli for all tasks included simple shapes and black and white clipart images, approximately 3.5cm x 3.5cm displayed on a black background (see Figures 2E, 3A, and 7A-B for examples of stimuli).

**Figure 3.**
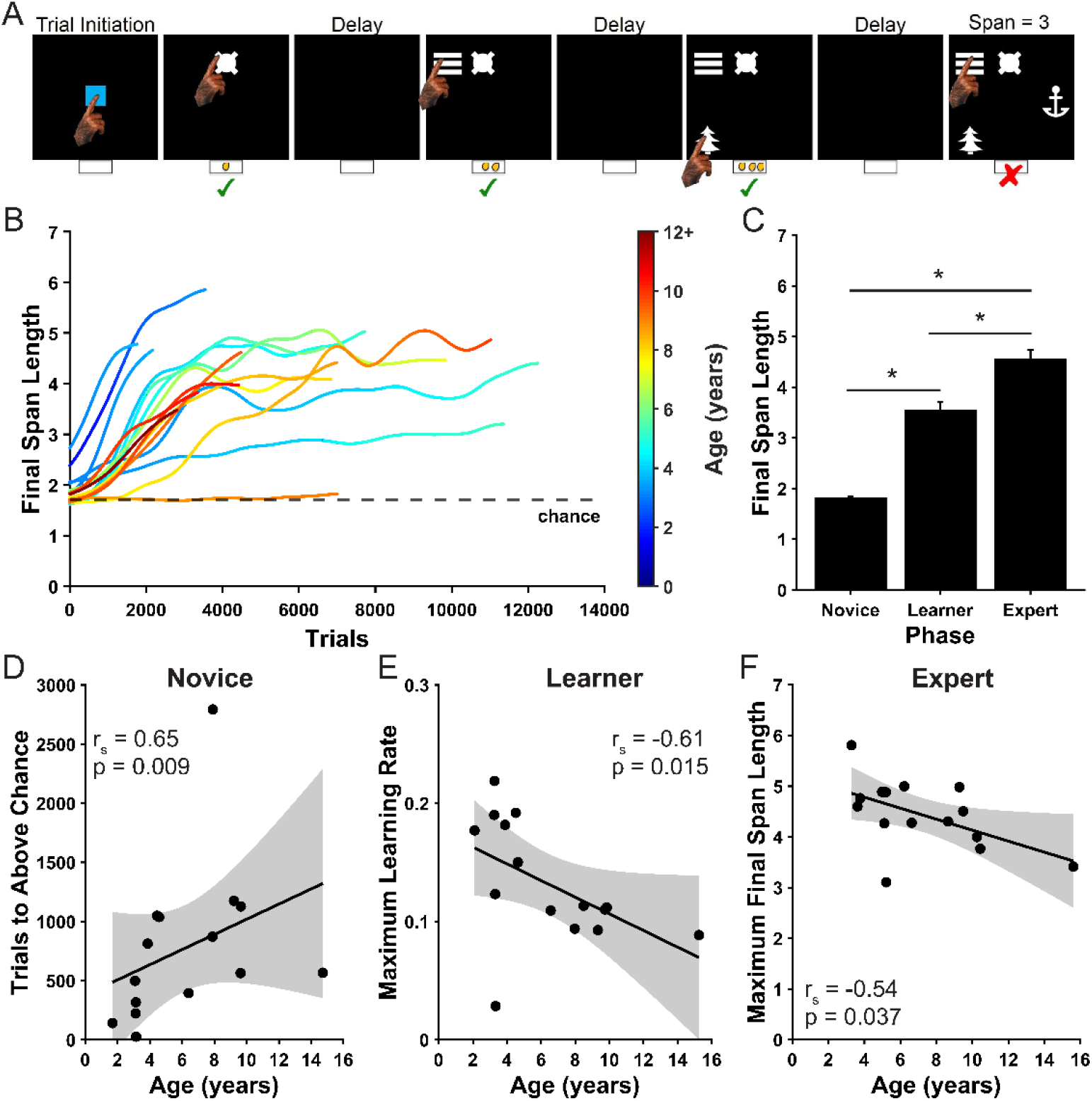
Delayed Recognition Span Task (DRST). A) Example of one DRST trial. Green check marks indicate correct responses. Red X indicates an error. B) Learning curves for each of the 16 marmosets trained to perform the task. Chance performance, determined with a Monte Carlo simulation, is indicated by the black dashed line. Colors of learning curves indicate the age of the marmoset, with color changing within each curve as the animal aged over the course of the experiment. C) Final Span Length increases with learning across the Novice (N), Learner (L), and Expert (E) Phases (mean ± SEM). Aging is correlated with D) increased duration of the Novice Phase, E) slowed learning rate during the Learning Phase, and F) decreased working memory abilities after achieving Expert status. Each black circle represents one animal. Linear regression models were used to determine 95% confidence intervals shaded in gray. X-axis in panel D is age at the end of the Novice Phase. X-axes in panels E and F correspond to age at the time of measurement. *p < 0.05.

**Figure 4.**
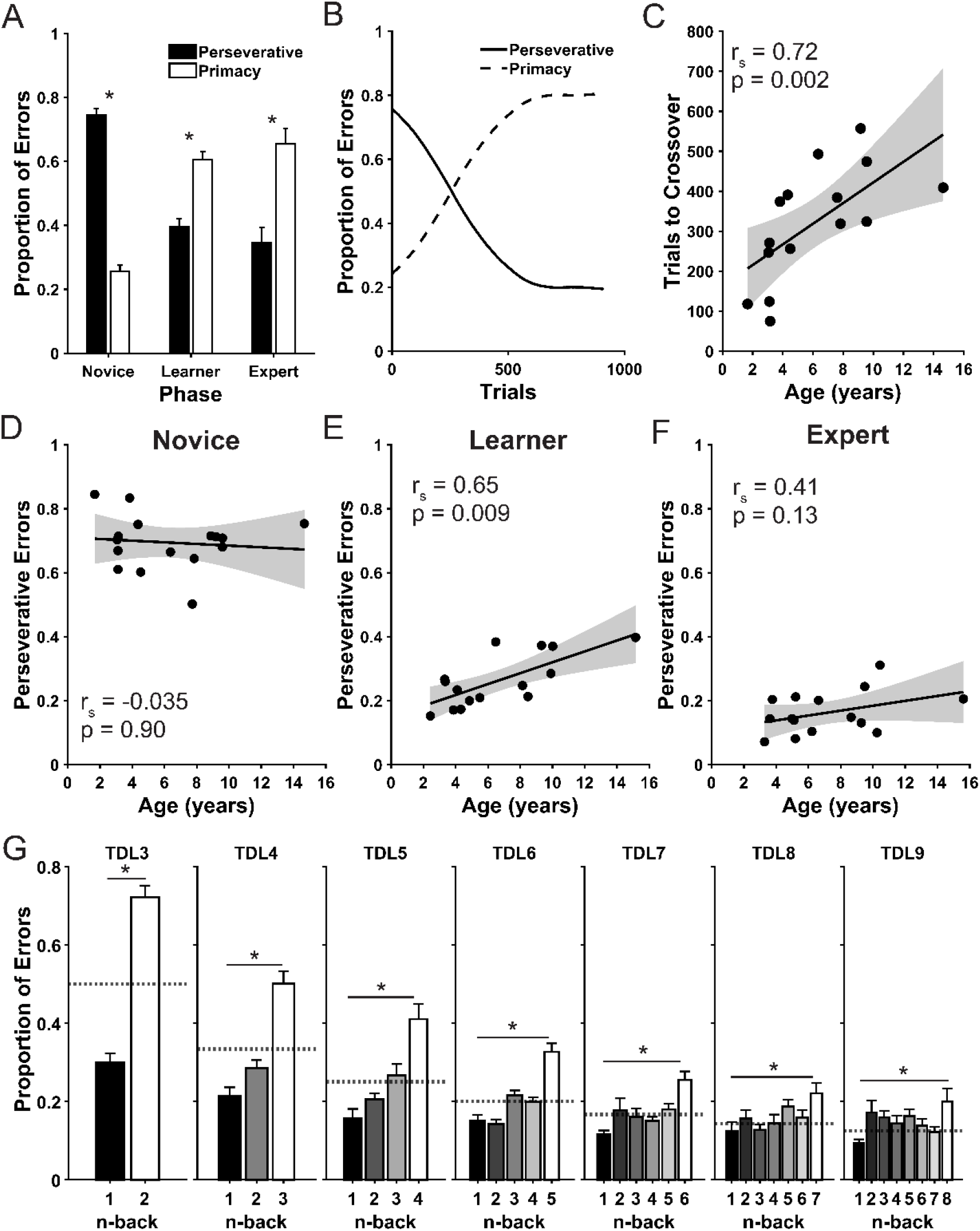
Error patterns. A) Errors transitioned from predominantly perseverative early in training to predominantly primacy errors as marmosets improved performance on the task. Bars are mean ± SEM. B) Representative example of the pattern of errors across trials in an individual marmoset. C) There was a strong significant correlation between age and the number of trials that occurred before the fraction of Primacy Errors exceeded the fraction of Perseverative Errors. Each black point corresponds to one animal. D-F) Correlations demonstrating that only during the Learner Phase is there a significant relationship between age and the proportion of errors that were perseverative across all TDLs combined. G) The distribution of errors made to each n-back on each TDL in the Expert Phase was significantly different from chance (dashed lines). Marmosets incorrectly identified remote (i.e., higher n-back) stimuli as novel more frequently than more recent stimuli (i.e., lower n-back). *p < 0.05.

**Figure 5.**
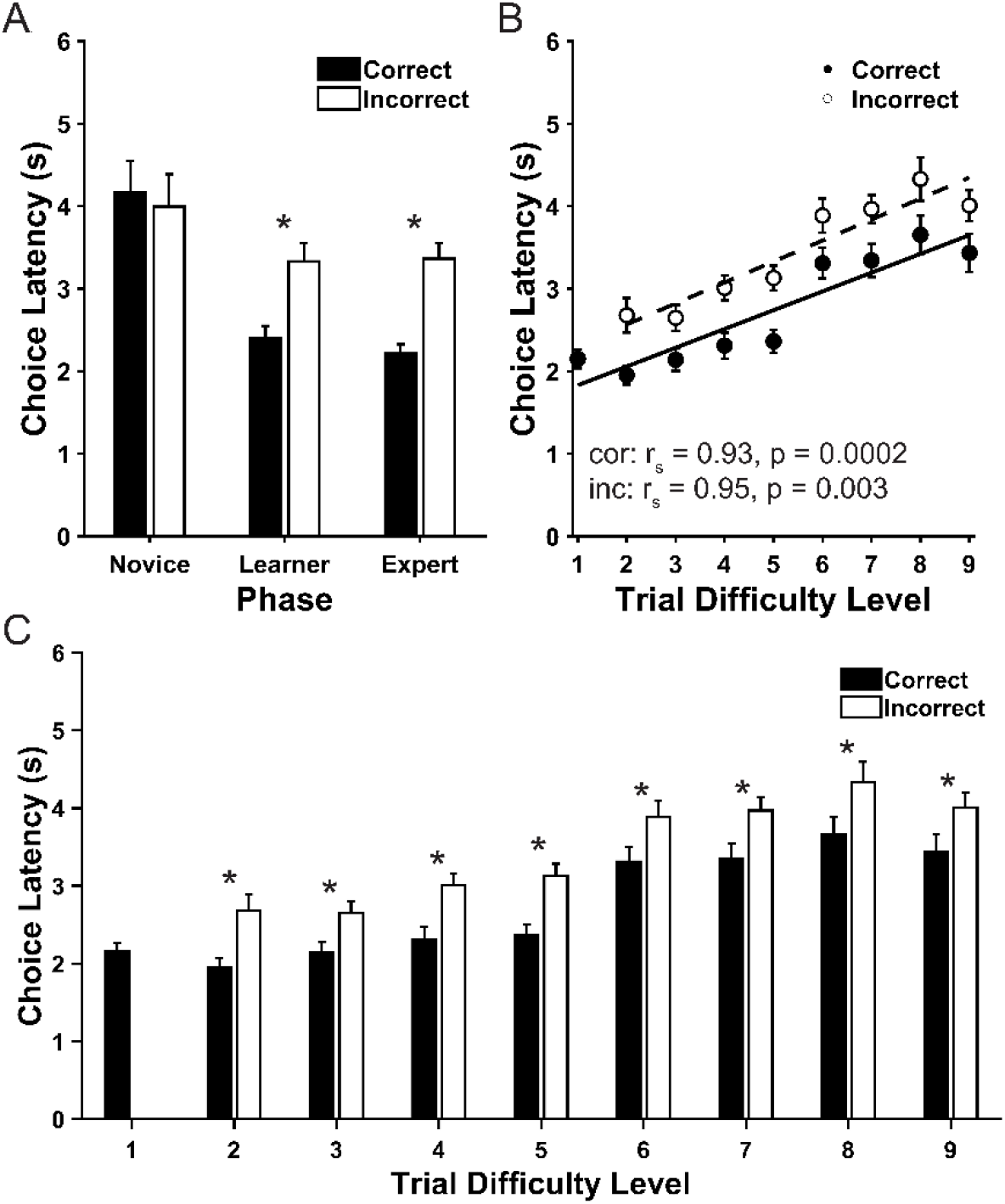
Choice latency patterns. A) Correct and incorrect choice latencies for each Phase of the DRST. Correct latencies decreased with task experience. Incorrect choice latencies were longer than correct choice latencies in the Learner and Expert Phases, showing errors were not due to impulsivity. B) There were strong positive correlations between trial difficulty level and incorrect or correct choice latencies demonstrating the increased cognitive load of more difficult trials. C) Incorrect choice latencies were longer than correct choice latencies for all trial difficulty levels, showing that, even when trials were the most difficult, errors were not due to impulsive choices. Black bars and circles are correct choice latency mean ± SEM. White bars and circles are incorrect choice latency mean ± SEM. *p < 0.05.

**Figure 6.**
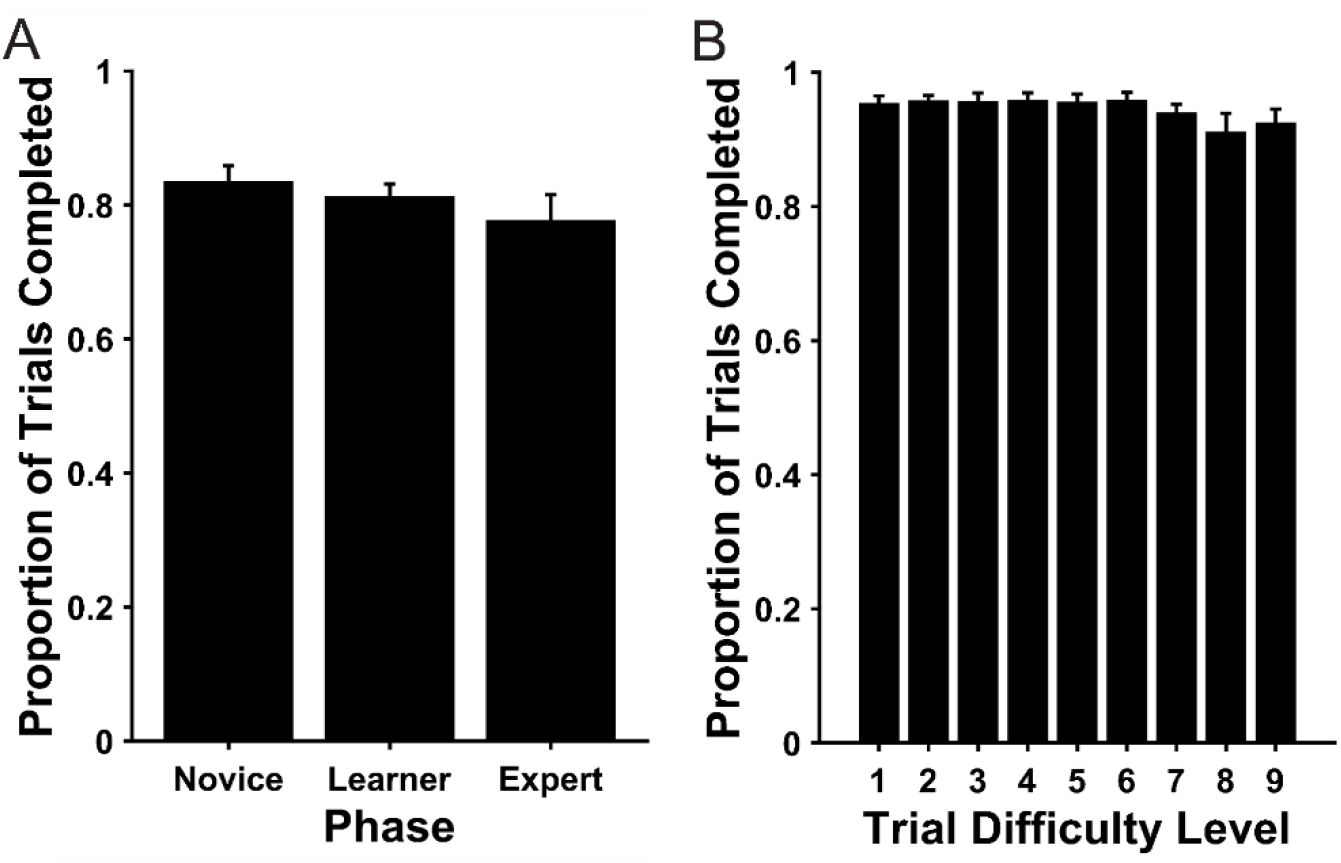
Trial completion rates. A) Marmosets completed a substantially large proportion of trials that they initiated across all Phases of the DRST. B) Continued engagement on even the most difficult portions of trials demonstrate that marmosets were highly motivated. Bars represent mean ± SEM.

**Figure 7.**
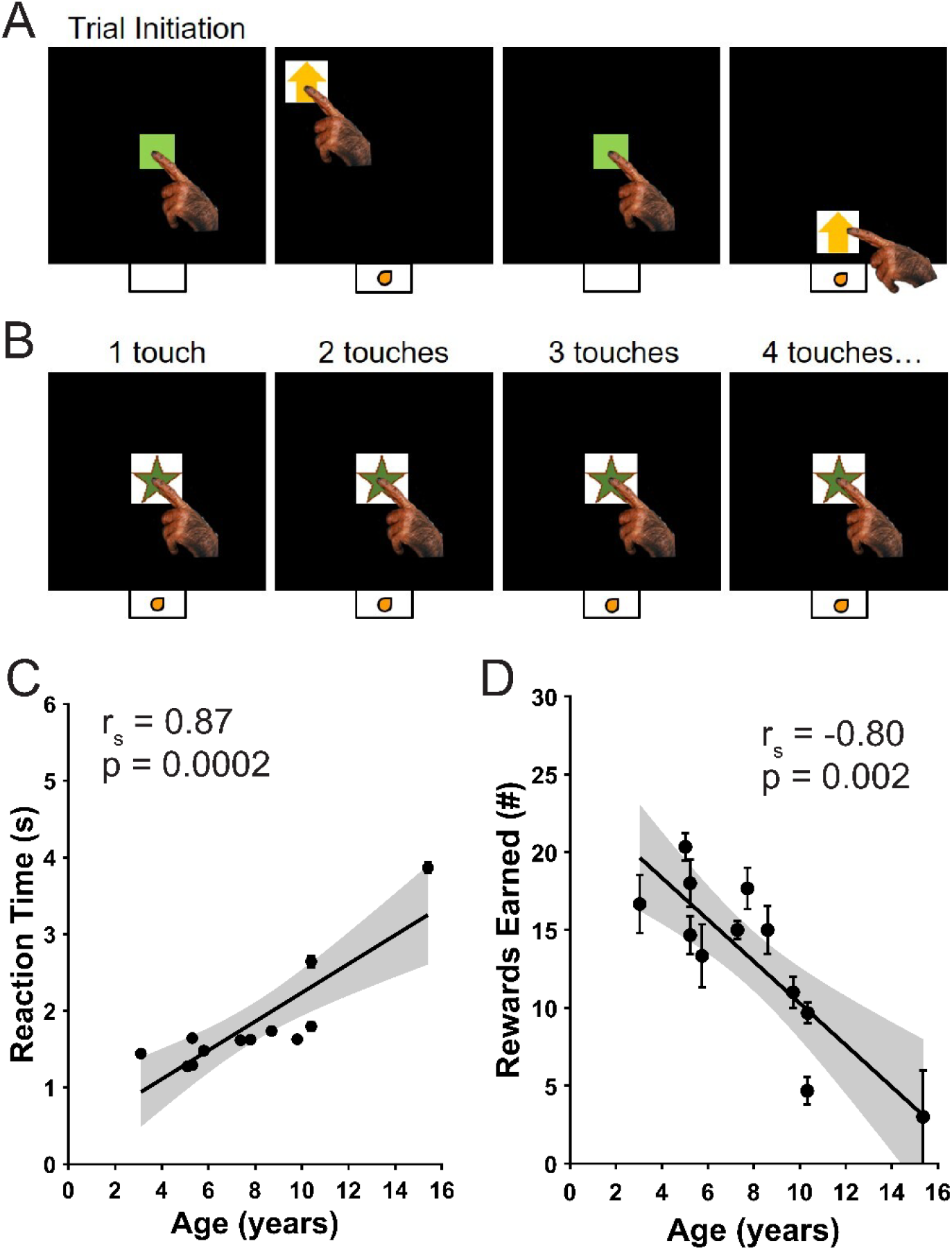
Non-cognitive control tasks. A) Example trials of the Reaction Time Task (RTT). B) Example trials of the Progressive Ratio Task (PRT). C) Data from the RTT showing a strong positive correlation between age and reaction time. Each plotted point represents the average ± SEM performance of one marmoset from 10 sessions of the RTT. D) Data from the PRT showing a strong negative correlation between age and number of rewards earned. Each plotted point represents the average ± SEM performance of one marmoset from three sessions of the PRT.

### Statistical Analyses

Data were analyzed using MATLAB (Mathworks, Natick, MA). Kolmogorov-Smirnov tests determined that the data were not normally distributed, and so non-parametric statistical analyses were used throughout the study. Therefore, correlations were assessed using Spearman’s rank-order correlations, Scheirer Ray Hare Tests were used to assess two factor interactions across time, and Friedman’s Tests with Nemenyi post-hoc tests, or Wilcoxon’s signed-rank tests, were used to identify within-factor differences. Performance was compared to chance using Chi-Square Goodness of Fit Tests. Chance levels were determined via a Monte Carlo simulation and p < 0.05 was considered significant.

### Cognitive Testing

#### Touch Training

Marmosets were naive to cognitive testing at the beginning of the study. Therefore, they were first trained to touch the screen to earn fluid rewards. The touch training process had three stages. The goals of the first stage were to habituate marmosets to the touch screen station, encourage physical engagement with the screen and the reward sink, and to associate touching the screen and reward delivery. To do this, a sweet substance (Marshmallow Fluff™) was applied on the screen, in small quantities, at each of the nine locations accessible through the mask. When the training session started, blue square target stimuli appeared in each of the nine locations, under the Fluff. When the marmosets touched the screen to obtain the Fluff a touch was detected and 0.2mL of reward was dispensed into the sink below the screen. The marmoset’s behavior was remotely observed via Ethernet-enabled cameras (RLK16-410B8, Reolink, Wilmington, DE). Animals were observed to identify when they learned that touching the screen caused a reward to be dispensed. Their behavior changed as they learned this association. Early in training the monkeys only consumed rewards when they noticed accumulation in the sink. When the association had been made, however, the monkey’s behavior changed, and immediately following each screen touch, they consumed the reward from the sink before touching the screen again. Once this association was established, the second stage of touch training began. The goals of the second stage were to characterize marmoset responses to each of the nine possible stimulus display locations, and to train the marmosets that only locations with stimuli have the potential to yield rewards, when selected. Initially, all nine locations had targets. As training continued over the course of approximately five daily training sessions, the number of target stimuli presented on the screen on each trial decreased in a stepwise manner from nine targets to one. During this stage, Fluff was no longer applied to the screen, and correct target selections were rewarded with 0.2mL of reward. Each target stimulus was placed in a randomly selected location. Once marmosets reliably touched the one randomly positioned target, the third and final stage of touch training commenced. The goal of the third stage was to maintain robust engagement with the task while reducing the volume of reward earned for each response. On each trial of this third stage, one stimulus was displayed on the screen in a randomly chosen location. Reward volumes were decreased from 0.2mL per response to a final volume of 0.05mL per response, in a stepwise manner over approximately four daily training sessions.

#### Delayed Recognition Span Task (Figure 3A)

The Delayed Recognition Span Task (DRST) measures working memory capacity. Each trial of the DRST was initiated by the marmoset touching a blue square stimulus in the center of the screen. Then, a single black and white stimulus, drawn randomly from a set of 400 images, was displayed on the screen in one of nine possible locations, also selected randomly. When the marmoset touched this first stimulus, a small liquid reward was dispensed. Following a two-second delay, during which the screen was blank, a two alternative forced choice was presented with the original stimulus appearing in its original location, and a second, novel stimulus appearing in a different pseudo randomly selected location. If the marmoset selected the novel object, a correct response was logged, a liquid reward was dispensed, and another two-second delay ensued. Following the delay, the first two stimuli appeared in their original locations, and a third, novel stimulus, also appeared, in a pseudo randomly chosen location. The marmoset was again rewarded for choosing the novel stimulus. Novel stimuli were added after subsequent delays until the trial was terminated in one of three ways: 1) the marmoset made nine correct selections in a row; 2) the marmoset failed to make a selection within 12 seconds (i.e., omitted); 3) the marmoset made an incorrect response (i.e., selected a non-novel stimulus). In the case of an omission or incorrect response, no reward was dispensed and a 5s time-out period began before the start of a new trial. For each trial, the number of stimuli correctly selected before the trial terminated was recorded as the “Final Span Length”. Reward volumes were increased with increased trial difficulty to encourage continued task engagement. Specifically, marmosets earned 0.05mL of reward for correct responses when one stimulus was on the screen, 0.1mL of reward for correct responses when two, three, or four stimuli were on the screen, and 0.2mL of reward for correct responses when five, six, seven, eight, or nine stimuli were on the screen. The number of objects on the screen is referred to below as trial difficulty levels (TDLs). Each marmoset was tested two to five days per week, and each testing session was terminated after three hours, or after the marmoset had earned 20mL of reward, whichever came first.

#### Reaction Time Task (Figure 7A)

The Reaction Time Task (RTT) measures the time required for the marmoset to reach out and touch the screen. It was used to distinguish age-related changes in non-cognitive motor speed from age-related changes in performance of the working memory task (DRST). Each trial of the RTT was initiated by the marmoset touching a green square stimulus in the center of the screen. Then, a target appeared in one of nine locations chosen pseudo-randomly. When the marmoset touched the target, they earned a 0.1mL liquid reward and the amount of time elapsed between target onset and selection by the marmoset was recorded as the reaction time on that trial. Marmosets were given one practice session to learn the task, and then data were collected from ten subsequent test sessions. Each of the 10 test sessions were concluded when marmosets had performed 108 trials or one hour had elapsed since the beginning of the testing session.

#### Progressive Ratio Task (Figure 7B)

The Progressive Ratio Task (PRT) measures motivation. It was used to distinguish age-related changes in non-cognitive motivation from age-related changes in DRST performance. Each trial of the PRT began with the presentation of a stimulus in the middle of the screen. Marmosets were rewarded for touching the stimulus under a progressive-ratio schedule of reinforcement where response requirements increased during a testing session. The initial response requirement was one touch to earn a reward. Once the monkey satisfied a response requirement, the stimulus was removed from the screen, reward dispensed, and then the stimulus was replaced on the screen. The response requirement increased by one until eight response requirements were completed, and subsequently doubled after every eight response requirements were completed thereafter. The specific response requirements used were: (increment = 1) 1, 2, 3, 4, 5, 6, 7, 8; (increment = 2) 10, 12, 14, 16, 18, 20, 22, 24; (increment =4) 28, 32, 36, 40, 44, 48, 52, 56; etc. Test sessions were concluded when one hour had elapsed or the marmoset reached a response requirement of 120 touches. Marmosets were given one practice session to learn the task, and then data were collected from three subsequent test sessions.

## RESULTS

### Delayed Recognition Span Task (DRST)

To assess whether marmosets demonstrate age-related working memory impairment, we implemented a touch screen version of the Delayed Recognition Span Task in a cohort of animals across a broad age range. In this task, marmosets initiated a trial and were then rewarded for touching a single visual stimulus (an “object”) appearing on the screen. Following a brief delay, during which the screen was blank, marmosets were rewarded for identifying the novel object in a two-alternative forced-choice. If correct, a brief delay was followed by the appearance of three objects; the two that had been seen earlier in the trial, and one novel. After each correct answer, reward was delivered, and an additional object was added to the array. When a marmoset chose a non-novel object, the trial ended without reward, and the trial sequence began anew.

The primary measure used to assess DRST performance was Final Span Length (FSL). As in prior work (Herndon et al., 1997; Moss et al., 1997; Killiany et al., 2000; Moore et al., 2017), this measure quantifies the animal’s working memory ability and is defined as the number of stimuli the marmoset correctly reported as novel on each trial. The raw data learning curves for each animal, plotted as a function of FSL over time, were smoothed using a Gaussian-weighted moving average over a 2000 trial window (Figure 3B), and a Monte Carlo simulation was used to model chance performance.

We analyzed performance during three distinct Phases along each marmoset’s learning curve, designated as “Novice”, “Learner”, and “Expert”. The Novice Phase was defined as the trials on which performance did not deviate from that expected by chance and began with the first trial ever performed. The end of the Novice Phase was determined using Kolmogorov-Smirnov Goodness of Fit Tests to identify the initial point where 100 successive experimentally measured FSL distributions differed significantly from a null distribution drawn from the Monte Carlo simulations. The Learner Phase began immediately after the Novice Phase and ended on the trial before the 90th percentile of performance. The Expert Phase encompassed trials with performance between the 90th and 100th percentiles, inclusive. These Phases were distinct from one another, with no trials falling into multiple categories. To validate these Phase definitions, we tested whether, as expected, average performance increased significantly across the three Phases and found that it did (Figure 3C; non-parametric Friedman’s test: χ^2^(2) = 30, p = 3.1×10^-7^); pairwise post-hoc Nemenyi tests: Novice vs Learner: p=0.017; Novice vs Expert: p = 1.4×10^-7^; Learner vs Expert: p = 0.017).

To determine if there is a relationship between age and the amount of experience necessary to acquire the rules of the DRST, we calculated the number of trials each marmoset completed before their performance significantly deviated from chance (i.e., the length of the Novice Phase). A Spearman’s rank-order correlation revealed a strong, positive association that was statistically significant, indicating that aging marmosets required more experience to initially acquire the DRST rules (Figure 3D; r_s_(13) = 0.65, p = 0.009). One marmoset failed to perform above chance levels, despite completing 7,000 DRST trials, and so this animal was excluded from the statistical analyses conducted on the Learner and Expert Phases, described below.

In the Learning Phase, marmosets demonstrated rapid improvement in performance. To determine if there were aging-associated differences in the rate of improvement in the Learning Phase, we assessed the maximum learning rate for each monkey. Maximum learning rate was calculated by finding the largest derivative of the mean FSL curve, computed over a rolling 100 trial window. A Spearman’s rank-order correlation revealed a significant negative correlation between age and maximum learning rate, demonstrating that with increasing age, learning rates slowed (Figure 3E; r_s_(13) = −0.61, p = 0.015).

In the Expert Phase, marmosets reached maximal levels of performance, and typically plateaued. We determined each marmoset’s maximum level of performance by calculating the mean FSL achieved during the Expert Phase. A Spearman’s rank-order correlation was used to assess the relationship between age and maximum FSL. It revealed a significant negative correlation between these variables, indicating that marmoset aging is associated with decreased working memory capacity (Figure 3F; r_s_(13) = −0.54, p = 0.037).

Additional information about cognitive performance and the way in which marmosets approached the task can be discerned by assessing patterns of errors made while performing the DRST. Instances when marmosets end a trial with a FSL of two (i.e., make an error on TDL3 when there are three objects on the screen), are especially informative since errors can only be made by selecting the first object of the trial (primacy error) or the most recently correct object of the trial (perseverative error). Prior studies of learning have found that animals have a natural tendency to continue to perform behaviors for which they have recently been rewarded, a behavioral pattern that can yield perseverative errors. Thus, we hypothesized that, prior to acquiring the rules of the DRST (i.e., in the Novice Phase), marmosets would make more perseverative errors relative to primacy errors, and that once the rules of the DRST had been successfully acquired (i.e., Learner and Expert Phases), marmosets would make more primacy errors relative to perseverative errors. To examine this we calculated the proportion of primacy and perseverative errors marmosets made on TDL3 trials during each of the three Learning Phases (Figure 4A) and found that there was a significant interaction between error type (perseverative, primacy) and Phase (Novice, Learner, Expert) (Scheirer Ray Hare Test: H(2) = 58.24, p = 2.0×10^-13^). Across Phases, there was a significant decrease in the fraction of errors that were perseverative and corresponding increase in the fraction of errors that were primacy (Friedman’s Test; χ^2^(2) = 22.93, p = 1.05×10^-5^). Specifically, this reached statistical significance between the Novice and Learner Phases and between the Novice and Expert Phases, but not between the Learner and Expert Phases (pairwise post-hoc Nemenyi tests; Novice vs Learner: p = 0.01; Novice vs Expert: p = 6.55×10^-6^; Learner vs Expert: p = 0.16). In line with our hypothesis, the predominant error type changed with learning of the task such that during the Novice Phase, errors were predominantly perseverative, whereas primacy errors were most prevalent in the Learner and Expert Phases (Wilcoxon signed-rank tests for paired samples; Novice: Z = 3.38, p = 6.10×10^-5^; Learner: Z = 2.641, p = 0.0054; Expert: Z = 2.527, p = 0.0084).

To determine if the slowed learning rates of aging marmosets could be explained by a protracted time course of the predominant error type change (“crossover”), we quantified the number of TDL3 trials before the crossover for each animal (Figures 4B-C). A Spearman’s rank-order correlation revealed a significant positive correlation between age and trials to the crossover, demonstrating that, on TDL3 trials, aging marmosets switch to predominantly primacy errors after more task experience than younger marmosets (Figure 4C; r_s_(13) = 0.72, p = 0.002).

Since the TDL3 error analyses revealed an age-related difference in the rate of error type switching from perseverative to primacy, we went on to examine if there was an age-related change in the magnitude of perseveration as well. Previous studies have found that aged macaque monkeys make more perseverative errors on the DRST than do young macaques (Moss et al., 1997). Therefore, we hypothesized that higher levels of perseveration could underlie the slowed learning rate of aging marmosets (Figure 3E) and may also explain the overall age-related working memory impairment (Figure 3F). To test this hypothesis, we calculated the proportion of perseverative errors across TDLs during each of the three Phases. We found that aging was associated with more perseverative errors, but only in the Learner Phase (Figures 4D-F; Spearman’s rank-order correlations; Novice: r_s_(14) = −0.035, p = 0.90; Learner: r_s_(13) = 0.65, p = 0.009; Expert: r_s_(13) = 0.411, p = 0.13). These results show that higher levels of perseveration underlie the slowed learning rate of aging marmosets, and that perseverative errors do not account for the age-related working memory impairment.

Trials of increased difficulty were reliably performed during the Expert Phase. The pattern of errors marmosets made on these trials provide an opportunity to examine whether marmosets succumb to working memory interference. To do this, we quantified the distribution of errors according to the distance in the past the incorrectly chosen stimulus was presented (i.e., “n-back”). For example, if a marmoset made an error when there were four objects on the screen (TDL4), they would earn a FSL of three on that trial. In this case, they could make an error by selecting the first object presented on the trial (n-back 3; primacy), the second object presented on the trial (n-back 2), or the third object presented on the trial (n-back 1; perseverative). The distribution of n-back errors made in the Expert Phase was quantified for each TDL and compared to chance performance using Chi-Square Goodness of Fit Tests (Figure 4H). These tests revealed that, for all TDLs, the observed n-back error distributions were significantly different from those expected by chance (TDL3: χ^2^ = 154.17, p = 2.13×10^-35^; TDL4: χ^2^ = 110.98, p = 7.94×10^-25^; TDL5: χ^2^ = 116.52, p = 4.34×10^-25^; TDL6: χ^2^ = 74.06, p = 3.15×10^-15^; TDL7: χ^2^ = 50.44, p = 1.13×10×10^-9^; TDL8: χ^2^ = 38.49, p = 9.01×10^-7^; TDL9: χ^2^ = 14.74, p = 0.04). Further, on each TDL, marmosets made significantly more primacy errors than perseverative errors (Wilcoxon signed-rank tests for paired samples; TDL3: Z = 3.38, p = 6.1×10^-5^; TDL4: Z = 3.05, p = 0.001; TDL5: Z = 3.24, p = 0.0002; TDL6: Z = 3.27, p = 0.0001; TDL7: Z = 3.15, p = 0.00024; TDL8: Z = 2.52, p = 0.0081; TDL9: Z = 2.10, p = 0.034). These results show that marmosets experienced retroactive interference, wherein newly acquired information disrupts temporary memory storage, leading to errors made by identification of more remotely presented stimuli as novel.

One advantage of using infrared touch screens to assess cognitive performance is the ability to measure choice latencies accurately and reliably. This measure is widely considered a robust readout of processing speed and is correlated with cognitive load and task difficulty (Bopp and Verhaeghen, 2018; Gray et al., 2018; De Boeck and Jeon, 2019). To determine if this pattern held true with marmosets performing the DRST, we assessed correct and incorrect choice latencies for each marmoset on each Phase of the DRST, and for each TDL in the Expert Phase. A Scheirer Ray Hare Test revealed significant main effects of latency type (correct choice, incorrect choice) and Phase of the DRST (Novice, Learner, Expert), and also a significant interaction between these factors (Figure 5A; latency type: H(1)=12.58, p=0.0004; Phase: H(2)=15.13, p=0.0005; interaction: H(2)=8.69, p=0.01). Correct choice latencies, but not incorrect choice latencies, significantly decreased across the Phases of the DRST (Friedman’s Tests; correct: χ^2^(2) = 22.93, p = 1.05×10^-5^; incorrect: χ^2^(2) = 0.12, p = 0.94). Correct choice latencies were shorter in the Learner and Expert Phases than the Novice Phase, with no difference between the Learner and Expert Phases (pairwise post-hoc Nemenyi tests; Novice vs Learner: p = 0.01; Novice vs Expert: p = 6.55×10^-5^; Learner vs Expert: p = 0.16). These results show that, with improvement on the task, marmosets were more quickly able to identify the novel stimulus. Next, to determine if incorrect choices could be due to impulsiveness, we compared correct to incorrect choice latencies within each of the Phases. During the Novice Phase, correct and incorrect choice latencies were similar, however during the Learner and Expert Phases, incorrect choice latencies were significantly longer than correct choice latencies (Figure 5A; Wilcoxon signed-rank tests for paired samples; Novice: Z = 0.71, p = 0.49; Learner and Expert: Z = 3.38, p = 6.10×10^-5^). This demonstrates that, when marmosets made an error, it was likely not due to impulsivity since they took considerably longer to respond in those cases.

To investigate choice latency patterns more thoroughly, we quantified the Expert Phase latency data by TDL. Since there was no significant relationship between age and choice latencies (Spearman’s rank-order correlations; correct: r_s_(13) = 0.40, p = 0.14; incorrect: r_s_(13) = 0.30, p = 0.27), further analyses focused on assessing whether the latency data revealed information about cognitive load and task difficulty. First, we examined if increased cognitive load required by more difficult TDLs was reflected in the choice latency data. Strong, positive Spearman’s rank-order correlations between both types of choice latency and trial difficulty level reflect longer processing time necessary for more difficult trials (Figure 5B; correct: r_s_(7) = 0.93, p = 0.0002; incorrect: r_s_(6) = 0.95, p = 0.0003). Finally, we assessed whether the overall pattern of longer incorrect than correct choice latencies found in the Expert Phase was consistent for each of the TDLs separately. A Scheirer Ray Hare Test revealed significant main effects of latency type (correct choice, incorrect choice; H(1) = 27.97, p = 1.2×10^-7^) and trial difficulty level (TDL1-9; H(7) = 94.12, p = 1.8×10^-17^), but no significant interaction (H(7) = 0.97, p = 1.00). Wilcoxon signed-rank tests for paired samples revealed incorrect choice latencies were longer than correct choice latencies at all TDLs, demonstrating that the marmosets took more time to make a choice if they made an error, even on the most difficult trials, and therefore were not choosing impulsively when incorrect (Figure 5C; TDL 2: Z = 3.26, p = 0.0001; TDL3: Z = 3.21, p = 0.0002; TDL4: Z = 3.26, p = 0.0001; TDL5: Z = 3.26, p = 0.0001; TDL6: Z = 3.20, p = 0.0002; TDL7: Z = 3.26, p = 0.0001; TDL8: Z = 3.14, p = 0.0004; TDL9: Z = 3.26, p = 0.0001).

To assess marmosets’ engagement on the DRST, we quantified the proportion of initiated trials that were completed versus omitted (no response within the 12 second response window). There was no significant difference in trial completion rate across the Novice, Learner, and Expert Phases (Figure 6A; Friedman’s Test; χ^2^(2) = 1.73, p = 0.42), indicating that marmosets were equivalently engaged with the task throughout all Phases. Further, there was no significant relationship between age and proportion of trials completed for any of the Phases (Novice: r_s_(14) = −0.044, p = 0.08; Learner: r_s_(14) = - 0.10, p = 0.73; Expert: r_s_(14) = −0.30, p = 0.27).

Finally, to assess whether marmosets were equally engaged on portions of trials that were more difficult (i.e., higher TDLs), we quantified completion rates for each of the TDLs independently. During the Expert phase, marmosets completed more than 90% of each of the TDLs they encountered, demonstrating continued engagement even when trials became challenging, despite not being water restricted, and testing in the home cage in a room that housed many other animals, and being free to leave the testing chamber at any time (Figure 6B).

### Reaction Time Task (RTT)

To assess potential non-cognitive confounds for interpretation of our DRST results, we measured motor speed using a Reaction Time Task. In this task, marmosets initiated a trial and then were rewarded for touching a single target stimulus placed pseudo randomly on the screen (Figure 7A). A Spearman’s rank-order correlation, used to determine the relationship between age and reaction time, revealed a strong, positive correlation between these variables, which reached statistical significance (Figure 7C; r_s_(10) = 0.87, p = 0.0002). This correlation was driven by specific strong associations between advancing age and longer reaction times on trials where the target stimulus appeared in the most difficult to reach locations on the screen:all three locations in the top row, and the two outer positions in the middle row (Top Row; Location 1: r_s_(10) = 0.72, p = 0.008; Location 2: r_s_(10) = 0.68, p = 0.015; Location 3: r_s_(10) = 0.70, p = 0.012; Middle Row; Location 4: r_s_(10) = 0.80, p = 0.002; Location 5: r_s_(10) = 0.54, p = 0.07; Location 6: r_s_(10) = 0.60, p = 0.04; Bottom Row; Location 7: r_s_(10) = 0.57, p = 0.054; Location 8: r_s_(10) = 0.44, p = 0.15; Location 9: r_s_(10) = 0.31, p = 0.32). These results demonstrate that with aging, marmosets have slowed motor speed.

To assess whether the age-related motor speed deficits were associated with DRST performance, we correlated reaction times on the RTT with several primary dependent variables from the DRST. There were no significant Spearman’s correlations between reaction time on the RTT and any of the DRST measures assessed including maximum learning rate (r_s_(10) = - 0.46, p = 0.14), maximal performance measured by FSL (r_s_(10) = −0.46, p = 0.13), and Expert Phase trial completion rate (r_s_(10) = −0.44, p = 0.15). These results show that even though marmosets have slowed motor speed with age, this did not negatively impact DRST performance.

### Progressive Ratio Task (PRT)

To assess another potential non-cognitive confound for interpretation of our DRST results, we measured motivation using a Progressive Ratio Task. In this task, marmosets were required to expend increasing amounts of effort (screen touches) to earn a same-sized reward (Figure 7B). A Spearman’s rank-order correlation was used to determine the relationship between age and the number of rewards earned during the PRT (Figure 7D). There was a strong, negative correlation between these variables, which was statistically significant, indicating that with advancing age, marmosets are less motivated to expend energy in exchange for a reward (r_s_(10) = −0.82, p = 0.001).

To assess the relationship between decreased motivation with age on the PRT and cognitive performance on the DRST, Spearman’s rank-order correlations were run between rewards earned on the PRT and the primary dependent variables from the DRST. There were no significant correlations between rewards earned on the PRT and any of the DRST measures assessed including maximum learning rate (r_s_(10) = 0.21, p = 0.52), Expert Phase FSL (r_s_(10) = 0.31, p = 0.33), and Expert Phase trial completion rate (r_s_(10) = 0.45, p = 0.14).

## DISCUSSION

This work firmly establishes the marmoset as a key model of age-related cognitive impairment with several advantages. This is important because, as an NHP, the marmoset offers translational relevance to humans beyond that found in rodents. As compared to the long-lived macaque, the short lifespan of marmosets enables longitudinal aging studies within a reasonably short timeframe. In this study, we performed comprehensive behavioral characterization of a large cohort of marmosets. These marmosets performed thousands of trials to assess learning and working memory capacity continuously for up to four years. This duration of continuous assessment accounts for up to 50 percent of the adult marmoset lifespan. Further, marmosets began training at ages that conjointly covered the lifespan, enabling assessment of age-related differences in acquisition of the working memory task. To our knowledge, these monkeys have undergone the most thorough cognitive profiling of any marmoset aging study conducted to-date, enabling us to identify which specific aspects of task performance are impaired with age.

### Comparison with prior studies of age-related cognitive changes in macaque and marmoset

Earlier DRST studies concluded that aged macaque monkeys have poorer working memory than young macaques (Herndon et al., 1997; Moss et al., 1997). The present results show that marmoset aging is associated with both learning and working memory impairments. Specifically, we found that aging is associated with delayed onset of learning, slowed learning rate after onset, and decreased asymptotic working memory performance. These impairments are evident in our study and may have been undetected in the macaque due to key differences in task implementation. In the macaque studies, monkeys were trained to a high level of proficiency on a delayed non-match-to-sample (DNMS) task prior to administration of a relatively small number of DRST trials (one hundred). This experimental design assumes that macaques can transfer the rule learned through DNMS training to DRST, with no additional training, and that they immediately perform the DRST maximally. It is unclear that they can, given the greater complexity of the DRST task. In contrast, the monkeys used in the present study trained on the DRST for thousands of trials over an extended time period. This enabled us to analyze both learning and maximal performance separately, using the same task. It is an open question whether the impaired performance of aged macaques reflects a memory impairment, a delayed onset of learning, a slower rate of learning, or some combination of these factors.

To-date, there is one prior longitudinal study of cognitive aging in marmosets (Rothwell et al., 2022). This prior study assessed visual discrimination and cognitive flexibility over a substantial portion of the adult marmoset lifespan. However, there are several key experimental design differences between this prior study and the work reported here, which merit discussion. First, Rothwell et al. tested marmosets once per year, whereas we measured cognitive performance continuously. The present approach densely characterized individual cognitive aging trajectories, facilitating analysis of multiple aspects of task performance. While we identified age-related impairment across all Phases of testing, Rothwell et al. reported impairment only on the last annual test. Second, marmosets enrolled in the Rothwell et al. study began testing at approximately the same age (4-6 years old), and impairment was identified when all animals were between 8-10 years of age. In contrast, marmosets in our study began testing at ages varying from young adult to geriatric, and we find that cognitive impairment occurs throughout the adult marmoset lifespan, in line with what has been observed in humans (Wu et al., 2021). Since aging is inherently a heterogeneous process, the fact that all marmosets in the Rothwell et al. study showed cognitive impairment simultaneously is unexpected. One possible explanation is that stimulus-dependent variation in task difficulty contributed to this pattern. For example, some stimulus pairs used in the first three years of the study were discriminable by shape and color, whereas all stimuli used in year four differed only by shape. Therefore, marmosets may have found the stimuli used in the terminal year more challenging, accounting for the decline in performance. Further, Rothwell et al. reported that impairment on the visual discrimination task was more severe than on the cognitive flexibility task. This is the opposite of what would be expected from published work demonstrating that cognitive flexibility often declines with aging, while visual discrimination typically does not (Bartus et al., 1979; Lai et al., 1995). Our results align with decades of work in macaques and humans, demonstrating the translational relevance of the marmoset as a model of aging.

### Patterns of errors reveal marmosets’ approach

Since detailed analysis of the types of errors made on neurocognitive assessments can, in humans, predict the emergence of future cognitive impairment (Bondi et al., 1999; Thomas et al., 2018), we investigated the patterns of errors marmosets made on the DRST as they learned the task and when they were performing maximally. Previous work in macaques has shown that aging is associated with increased perseverative errors on the DRST (Moss et al., 1997). Interestingly, our results only recapitulated this finding during the Learning Phase, and not during the Novice or Expert Phases, supporting the idea that aged macaque DRST impairment reflects learning rather than working memory deficits. In addition, older marmosets required more experience before they switched from mostly perseverative to mostly primacy errors. We hypothesize that both findings are associated with the learning impairment we identified in marmosets and supports learning rather than working memory impairment in the aged macaques. Finally, we used the pattern of error choices to determine whether marmoset performance was affected more significantly by proactive interference (i.e., full working memory prohibits adding novel objects), or retroactive interference (i.e., newly encountered information obscures more remote memories). We found that, once marmosets were performing maximally, they more frequently made an error by selecting stimuli that appeared earlier in the trial. This supports the idea that marmosets had more difficulty remembering stimuli that were more remote, and that performance was affected most significantly by retroactive interference (Amit et al., 2003). Together, these findings highlight the importance of analyzing process scores in addition to performance scores to understand behavior more thoroughly (Kaplan, 1988).

### Age-related motor speed deficits

Age-related motor speed deficits, measured by increased reaction times, are well-documented in rodents, NHPs, and humans (Bachevalier et al., 1991; Woods et al., 2015). To understand the impact of age on reaction time in our marmosets, and to assist us in controlling for this non-cognitive confound when interpreting our DRST results, we employed the RTT in a subset of the marmosets that performed the DRST. We found a strong association between increased reaction times and increased age. Critically, we found no associations between reaction time on the RTT and any of the DRST performance scores, demonstrating that the age-related motor speed impairments do not account for impairments on the DRST.

Unlike our findings from the RTT, we did not find any associations between aging and choice latencies on the DRST. We did, however, find that choice latencies were longer for more difficult TDLs, in all animals. This is likely because choice latencies on cognitive tasks are approximations of task difficulty and cognitive load (Bopp and Verhaeghen, 2018), and this supersedes the effects of age-related motor deficits. Additionally, for all TDLs, marmosets responded more slowly if their answer was incorrect than correct, demonstrating that, even when trials became more difficult, marmosets did not respond impulsively.

### Age-related motivation deficits

It is well-known that older rodents, NHPs, and humans have decreased motivation to earn rewards (Lanctôt et al., 2017; Lizarraga et al., 2020; Jackson et al., 2021). To evaluate the potential impact of age-related changes in motivation on DRST performance, we employed the well-established PRT to quantify motivation in our marmosets (Spinelli et al., 2004). We found a strong association between increasing age and decreasing motivation to earn rewards. We did not, however, identify any relationship between motivation on the PRT and any of the DRST performance scores, demonstrating that age-related decreases in motivation did not account for impairments on the DRST.

To directly assess motivation to perform the DRST, we quantified the proportion of trials that monkeys completed versus omitted. We found no significant relationship between age and proportion of trials omitted, unlike what we would have predicted from marmosets’ performance on the PRT. Furthermore, marmosets were equally likely to complete trials of each TDL, demonstrating engagement on the DRST even when trials were difficult. We hypothesize that when marmosets failed to complete a trial, it was likely due to inattention or distraction rather than low motivation.

### Summary

Here we have established the marmoset as a key model of age-related cognitive impairment. Our approach densely characterized cognitive aging trajectories in a large cohort of marmosets performing a translationally relevant working memory task. Through rigorous evaluation of process scores, we show that cognitive impairment can be detected long before it can be measured by overall performance scores. This framework will lead to better understanding of the aging process itself and may reveal why it is the biggest risk factor for neurodegenerative diseases.

## Acknowledgments

This research was supported by an AHA-Allen Initiative in Brain Health and Cognitive Impairment award made jointly through the American Heart Association and The Paul G. Allen Frontiers Group: 19PABH134610000AHA, a National Institutes of Health grant 1R21AG068967-01, grants from the Larry L. Hillblom Foundation and the Don and Lorraine Freeberg Foundation, and the Fiona and Sanjay Jha Chair in Neuroscience. We thank Katie Williams for assistance in the care of the marmosets and technical support.

## REFERENCES

Abbott DH, Barnett DK, Colman RJ, Yamamoto ME, Schultz-Darken NJ (2003) Aspects of common marmoset basic biology and life history important for biomedical research. Comp Med 53:339–350.

Amit DJ, Bernacchia A, Yakovlev V (2003) Multiple-object working memory--a model for behavioral performance. Cereb Cortex 13:435–443.

Bachevalier J, Landis LS, Walker LC, Brickson M, Mishkin M, Price DL, Cork LC (1991) Aged monkeys exhibit behavioral deficits indicative of widespread cerebral dysfunction. Neurobiology of Aging 12:99–111.

Bartus RT, Dean RL, Fleming DL (1979) Aging in the rhesus monkey: effects on visual discrimination learning and reversal learning. J Gerontol 34:209–219.

Beason-Held LL, Rosene DL, Killiany RJ, Moss MB (1999) Hippocampal formation lesions produce memory impairment in the rhesus monkey. Hippocampus 9:562–574.

Belham FS, Satler C, Garcia A, Tomaz C, Gasbarri A, Rego A, Tavares MCH (2013) Age-Related Differences in Cortical Activity during a Visuo-Spatial Working Memory Task with Facial Stimuli. PLoS One 8:e75778.

Belleville S, Rouleau N, Caza N (1998) Effect of normal aging on the manipulation of information in working memory. Mem Cogn 26:572–583.

Bondi MW, Salmon DP, Galasko D, Thomas RG, Thal LJ (1999) Neuropsychological function and apolipoprotein E genotype in the preclinical detection of Alzheimer’s disease. Psychol Aging 14:295–303.

Bopp KL, Verhaeghen P (2018) Aging and n-Back Performance: A Meta-Analysis. The Journals of Gerontology: Series B Available at: https://academic.oup.com/psychsocgerontology/advance-article/doi/10.1093/geronb/gby024/4944520 [Accessed May 18, 2022].

Bor D, Duncan J, Lee ACH, Parr A, Owen AM (2006) Frontal lobe involvement in spatial span: Converging studies of normal and impaired function. Neuropsychologia 44:229–237.

Cappell KA, Gmeindl L, Reuter-Lorenz PA (2010) Age differences in prefontal recruitment during verbal working memory maintenance depend on memory load. Cortex 46:462–473.

Comrie AE, Gray DT, Smith AC, Barnes CA (2018) Different macaque models of cognitive aging exhibit task-dependent behavioral disparities. Behav Brain Res 344:110–119.

De Boeck P, Jeon M (2019) An Overview of Models for Response Times and Processes in Cognitive Tests. Front Psychol 10:102.

De Castro V, Girard P (2021) Location and temporal memory of objects declines in aged marmosets (Callithrix jacchus). Scientific Reports 11.

Gazzaley A, Cooney JW, Rissman J, D’Esposito M (2005) Top-down suppression deficit underlies working memory impairment in normal aging. Nat Neurosci 8:1298–1300.

Glisky EL (2007) Changes in Cognitive Function in Human Aging. In: Brain Aging: Models, Methods, and Mechanisms (Riddle DR, ed) Frontiers in Neuroscience. Boca Raton (FL): CRC Press/Taylor & Francis. Available at: http://www.ncbi.nlm.nih.gov/books/NBK3885/ [Accessed May 18, 2022].

Gray DT, Umapathy L, Burke SN, Trouard TP, Barnes CA (2018) Tract-Specific White Matter Correlates of Age-Related Reward Devaluation Deficits in Macaque Monkeys. J Neuroimaging Psychiatry Neurol 3:13–26.

Herndon JG, Moss MB, Rosene DL, Killiany RJ (1997) Patterns of cognitive decline in aged rhesus monkeys. Behavioural Brain Research 87:25–34.

Hofer SM, Sliwinski MJ (2001) Understanding Ageing. An evaluation of research designs for assessing the interdependence of ageing-related changes. Gerontology 47:341–352.

Izpisua Belmonte JC et al. (2015) Brains, genes, and primates. Neuron 86:617–631.

Jackson MG, Lightman SL, Gilmour G, Marston H, Robinson ESJ (2021) Evidence for deficits in behavioural and physiological responses in aged mice relevant to the psychiatric symptom of apathy. Brain Neurosci Adv 5:23982128211015110.

Jeneson A, Mauldin KN, Squire LR (2010) Intact working memory for relational information after medial temporal lobe damage. The Journal of neuroscience: the official journal of the Society for Neuroscience 30:13624–13629.

Kaplan E (1988) The process approach to neuropsychological assessment. Aphasiology 2:309–311.

Killiany RJ, Moss MB, Rosene DL, Herndon J (2000) Recognition memory function in early senescent rhesus monkeys. Psychobiology 28:45–56.

Klencklen G, Banta Lavenex P, Brandner C, Lavenex P (2017) Working memory decline in normal aging: Memory load and representational demands affect performance. Learning and Motivation 60:10–22.

Lai ZC, Moss MB, Killiany RJ, Rosene DL, Herndon JG (1995) Executive system dysfunction in the aged monkey: Spatial and object reversal learning. Neurobiology of Aging 16:947–954.

Lanctôt KL, Agüera-Ortiz L, Brodaty H, Francis PT, Geda YE, Ismail Z, Marshall GA, Mortby ME, Onyike CU, Padala PR, Politis AM, Rosenberg PB, Siegel E, Sultzer DL, Abraham EH (2017) Apathy associated with neurocognitive disorders: Recent progress and future directions. Alzheimers Dement 13:84–100.

Lizarraga S, Daadi EW, Roy-Choudhury G, Daadi MM (2020) Age-related cognitive decline in baboons: modeling the prodromal phase of Alzheimer’s disease and related dementias. Aging (Albany NY) 12:10099–10116.

Maylor EA, Simpson EE, Secker DL, Meunier N, Andriollo-Sanchez M, Polito A, Stewart-Knox B, Mcconville C, O’connor JM, Coudray C (2006) Effects of zinc supplementation on cognitive function in healthy middle-aged and older adults: the ZENITH study. British Journal of Nutrition 96:752–760.

Mazurek A, Bhoopathy R, Read JCA, Gallagher P, Smulders TV (2015) Effects of age on a real-world What-Where-When memory task. Frontiers in Aging Neuroscience 7:1–17.

McQuail JA, Dunn AR, Stern Y, Barnes CA, Kempermann G, Rapp PR, Kaczorowski CC, Foster TC (2021) Cognitive Reserve in Model Systems for Mechanistic Discovery: The Importance of Longitudinal Studies. Front Aging Neurosci 12:607685.

Moore TL, Bowley B, Shultz P, Calderazzo S, Shobin E, Killiany RJ, Rosene DL, Moss MB (2017) Chronic curcumin treatment improves spatial working memory but not recognition memory in middle-aged rhesus monkeys. GeroScience 39:571–584.

Morrison JH, Baxter MG (2012) The ageing cortical synapse: Hallmarks and implications for cognitive decline. Nature Reviews Neuroscience 13:240–250.

Moss MB, Killiany RJ, Lai ZC, Rosene DL, Herndon JG (1997) Recognition memory span in rhesus monkeys of advanced age. Neurobiology of Aging 18:13–19.

Moss MB, Moore TL, Schettler SP, Killiany R, Rosene D (2007) Successful vs. Unsuccessful Aging in the Rhesus Monkey. In: Brain Aging: Models, Methods, and Mechanisms (Riddle DR, ed) Frontiers in Neuroscience. Boca Raton (FL): CRC Press/Taylor & Francis. Available at: http://www.ncbi.nlm.nih.gov/books/NBK1833/ [Accessed May 19, 2022].

Munger EL, Takemoto A, Raghanti MA, Nakamura K (2017) Visual discrimination and reversal learning in aged common marmosets (Callithrix jacchus). Neuroscience research 124:57–62.

Nyberg L, Boraxbekk C-J, Sörman DE, Hansson P, Herlitz A, Kauppi K, Ljungberg JK, Lövheim H, Lundquist A, Adolfsson AN, Oudin A, Pudas S, Rönnlund M, Stiernstedt M, Sundström A, Adolfsson R (2020) Biological and environmental predictors of heterogeneity in neurocognitive ageing: Evidence from Betula and other longitudinal studies. Ageing Res Rev 64:101184.

Phillips KA, Watson CM, Bearman A, Knippenberg AR, Adams J, Ross C, Tardif SD (2019) Age-related changes in myelin of axons of the corpus callosum and cognitive decline in common marmosets. American Journal of Primatology 81:e22949.

Preuss TM (1995) Do rats have prefrontal cortex? The rose-woolsey-akert program reconsidered. J Cogn Neurosci 7:1–24.

Rapp PR, Amaral DG (1989) Evidence for task-dependent memory dysfunction in the aged monkey. J Neurosci 9:3568–3576.

Rothwell ES, Workman KP, Wang D, Lacreuse A (2022) Sex differences in cognitive aging: a 4-year longitudinal study in marmosets. Neurobiol Aging 109:88–99.

Rusinek H, De Santi S, Frid D, Tsui W-H, Tarshish CY, Convit A, de Leon MJ (2003) Regional Brain Atrophy Rate Predicts Future Cognitive Decline: 6-year Longitudinal MR Imaging Study of Normal Aging. Radiology 229:691–696.

Sadoun A, Rosito M, Fonta C, Girard P (2019) Key periods of cognitive decline in a nonhuman primate model of cognitive aging, the common marmoset (Callithrix jacchus). Neurobiology of Aging 74:1–14.

Satler C, Belham FS, Garcia A, Tomaz C, Tavares MCH (2015) Computerized spatial delayed recognition span task: A specific tool to assess visuospatial working memory. Frontiers in Aging Neuroscience 7:1–9.

Schultz-Darken NJ, Braun KM, Emborg ME (2016) Neurobehavioral development of common marmoset monkeys. Developmental Psychobiology 58:141–158.

Spinelli S, Pennanen L, Dettling AC, Feldon J, Higgins GA, Pryce CR (2004) Performance of the marmoset monkey on computerized tasks of attention and working memory. Cognitive Brain Research 19:123–137.

Tardif SD (2019) Marmosets as a translational aging model—Introduction. American Journal of Primatology 81:1–3.

Thomas KR, Edmonds EC, Eppig J, Salmon DP, Bondi MW, Alzheimer’s Disease Neuroimaging Initiative (2018) Using Neuropsychological Process Scores to Identify Subtle Cognitive Decline and Predict Progression to Mild Cognitive Impairment. J Alzheimers Dis 64:195–204.

Upright NA, Baxter MG (2021) Prefrontal cortex and cognitive aging in macaque monkeys. Am J Primatol 83:e23250.

Voytko ML, Tinkler GP (2004) Cognitive function and its neural mechanisms in nonhuman primate models of aging, Alzheimer’s disease, and menopause. Frontiers in Bioscience 9:1899–1914.

Wise SP (2008) Forward frontal fields: phylogeny and fundamental function. Trends Neurosci 31:599–608.

Woods DL, Wyma JM, Yund EW, Herron TJ, Reed B (2015) Age-related slowing of response selection and production in a visual choice reaction time task. Front Hum Neurosci 9:193.

Wu Z, Woods RL, Wolfe R, Storey E, Chong TTJ, Shah RC, Orchard SG, McNeil JJ, Murray AM, Ryan J (2021) Trajectories of cognitive function in community-dwelling older adults: A longitudinal study of population heterogeneity. Alzheimers Dement (Amst) 13:e12180.

